# Formulas for Determining the Recovery Times for Optimal Two-Point Saturation–Recovery Measurement of the MR Longitudinal Relaxation Rate

**DOI:** 10.1101/534891

**Authors:** Jung-Jiin Hsu

**Author notes:** Also known as Jason Hsu.

## Abstract

The saturation–recovery method is a frequently used NMR/MRI technique for the measurement of the longitudinal relaxation rate *R*_1_. Under noise influence, the accuracy and the precision of the measurement outcome depend on the selection of the recovery times. This work seeks to determine the optimal recovery-time selection when the total scan time is constrained. A Monte Carlo computational method was used to simulate the noise-influenced distribution of the *R*_1_ measurement for various combinations of the two recovery times and to find the combination that produces the minimal standard deviation. Using the sum of recovery times (SRT) as the total scan time, mathematical formulas are derived from the simulation covering the SRT range of [0.05*T*_1_, 15*T*_1_]. The formulas describe the relationship between the SRT, the optimal accuracy and precision, and the recovery-time selection, and reproduce the results for the cases previously validated with experiments. In clinical application, the scan time is limited by the scan subject’s tolerance in the scanner and the time that can be allocated to the measurement in a scan session. These formulas provide a practical way to evaluate the trade-offs between precision and the scan time in scan planning.

## INTRODUCTION

Measuring the MR longitudinal relaxation rate *R*_1_ is a common procedure in NMR studies and in quantitative MRI. The saturation–recovery (SR) method is a popular technique for the *R*_1_ measurement, in which the longitudinal magnetization is set to zero by a 90° rf pulse and then measured after a delay time *τ* by recording the NMR signal; the delay is referred to as the recovery time and is a major variable that determines the time required for the measurement. The magnetization measurement is repeated for at least two different recovery times and then the value of *R*_1_ is determined from these measurements. A practical question is how the recovery times should be selected and how the selection affects the accuracy of the measurement. In clinical application, the imaging time is limited for patient comfort and by the available time. Therefore, the selection of the recovery times is further subject to a time constraint which is usually predetermined. This work presents practical formulas for determining such recovery times and for estimating the accuracy and precision of the measurement outcome for the SR method of two recovery times (i.e., two points).

In addition to the recovery times, time is also required for generating the image data, such as for playing out the RF and the encoding field gradient pulses, which can be significantly long in high-resolution imaging. The two-point SR method is thus of particular interest because it is the minimum requirement of the data to be acquired, and, therefore, often the first to be evaluated in developing a clinical imaging protocol.

An analytical calculus approach, e.g., Refs. [1–7], to these optimization questions would be very involved because of the increased mathematical complexity when the total imaging time is further subject to a constraint. Alternatively, a Monte Carlo computational study [8] was performed to answer these questions and the results were demonstrated with human subjects successfully. The Monte Carlo study was performed for a few settings of the total imaging time meant for experimental validation of the methodology. To facilitate quantitative MRI in clinical application, a systematic Monte Carlo computation for the two-point SR method is carried out in this work to derive general formulas covering a wide range of the total imaging time. These formulas describe the optimal selection of the two recovery times as a function of the total imaging time and the relationship between the total imaging time and the best expected measurement accuracy and precision.

## METHODS

The SR equation is given by

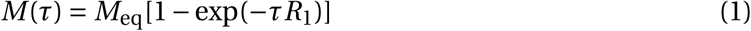

where *M* (*τ*) and *M*_eq_ represent the magnetization at recovery time *τ* and at the thermal equilibrium, respectively. Given two data points, *M* (*τ*_1_) and *M* (*τ*_2_), the SR equation can be solved to obtain *R*_1_ [8]. In this work, *T*_1_ represents the true longitudinal relaxation time of the specimen and is used as the unit of time; 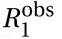 is the longitudinal relaxation rate derived from data and is expressed in units of 1/*T*_1_. The time constraint on the total imaging time is specified in terms of the sum of the recovery times (SRT) *T*_Σ_ = *τ*_1_ + *τ*_2_. The SRT is a measure of the total imaging time.

The range of the SRT considered is 0.05*T*_1_ ≤ *T*_Σ_ ≤ 15*T*_1_ and is represented by samples equally spaced by 0.25*T*_1_. For each sample, the optimal *τ*_1_, and hence *τ*_2_ = *T*_Σ_ -*τ*_1_, is determined by the Monte Carlo method described in the next subsection. A mathematical formula as a function of SRT is then derived by least-squares fitting the optimal *τ*_1_’s of all SRT samples to an empirical equation.

### Computational

For a given selection of the recovery times, the values of *R*_1_ derived from all possible results of the noise-influenced magnetization measurement form a distribution. The distribution is characterized by the signal-to-noise ratio (SNR), the SRT, and *τ*_1_. The distribution was simulated by using a large number of computer-synthesized data *M* (*τ*_1_) and *M* (*τ*_2_). Fourteen samples of SNR in [50, 64000] were considered, including 50, 80, 110, 140, 190, 240, 500, 1000, 2000, 4000, 8000, 16000, 32000, and 64000. The data for the measurement of *M* (*τ*) was synthesized as the modulus of a complex-valued number, in which the real part was calculated by the SR equation, Eq. (1), plus a computer-generated random number and the imaginary part was only a separately-generated random number. The SNR *ρ* is defined as [8]

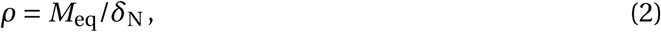

where *δ*_N_ is a representative noise amplitude. In an image, *δ*_N_ can be estimated by the average pixel intensity of a region of background where the signal is supposed to be zero. In the Monte Carlo computation, the noise in the image is modeled by the Rayleigh distribution and is simulated by a zero-mean normally-distributed random number generator of standard deviation *σ*, where

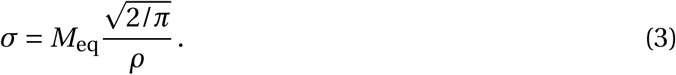

Because the signal amplitude *M* varies as a function of the recovery time *τ* as described by Eq. (1), the definition of SNR uses *M*_eq_ instead of *M* to characterize the SNR of the imaging environment rather than in the individual images [8].

For each sample of the SRT and SNR, an *R*_1_ distribution (consisted of 65536 trials) was established for each lattice point on a two-dimensional Cartesian grid; the two dimensions were *τ*_1_ and *τ*_2_ with grid spacing of 0.05*T*_1_. Because of the symmetry, only lattice points of *τ*_1_ < *τ*_2_ were considered.

For each sample of the SRT and SNR, the grid was searched for the lattice point whose *R*_1_ distribution has the smallest SD, and then the SD together with those of the nearest neighbors on the grid (if available) were fitted to a second order polynomial of *τ*_1_; from this polynomial, the true minimum SD and the optimal *τ*_1_ can be better located, which might be between the lattice points of the grid.

The search for the optimal *τ*_1_ was repeated for all the SRT samples in the interval 0.05*T*_1_ ≤ *T*_Σ_ ≤ 15*T*_1_ to establish the optimal recovery times as functions of the SRT. The computer programs in this work were coded by the present author in the C language. The routines in the GNU Scientific Library (http://www.gnu.org/software/gsl/) were used to implement the random-number generator.

## RESULTS

### Optimal Recovery Times vs. SRT

Figure 1 shows the optimal *τ*_1_ for a few sample SNR levels and the curve determined by the least-squares fitting. Because the optimal *τ*_1_ apparently does not depend on the SNR as shown in the figure, the curve fitting includes data of all SNR levels considered in this work and the resultant curve can be described by

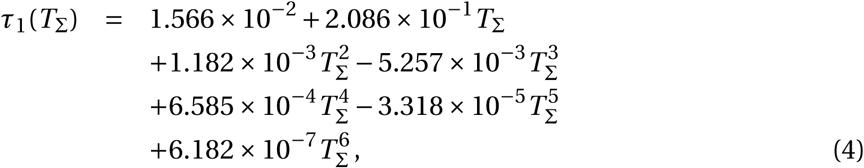

**Figure 1.**
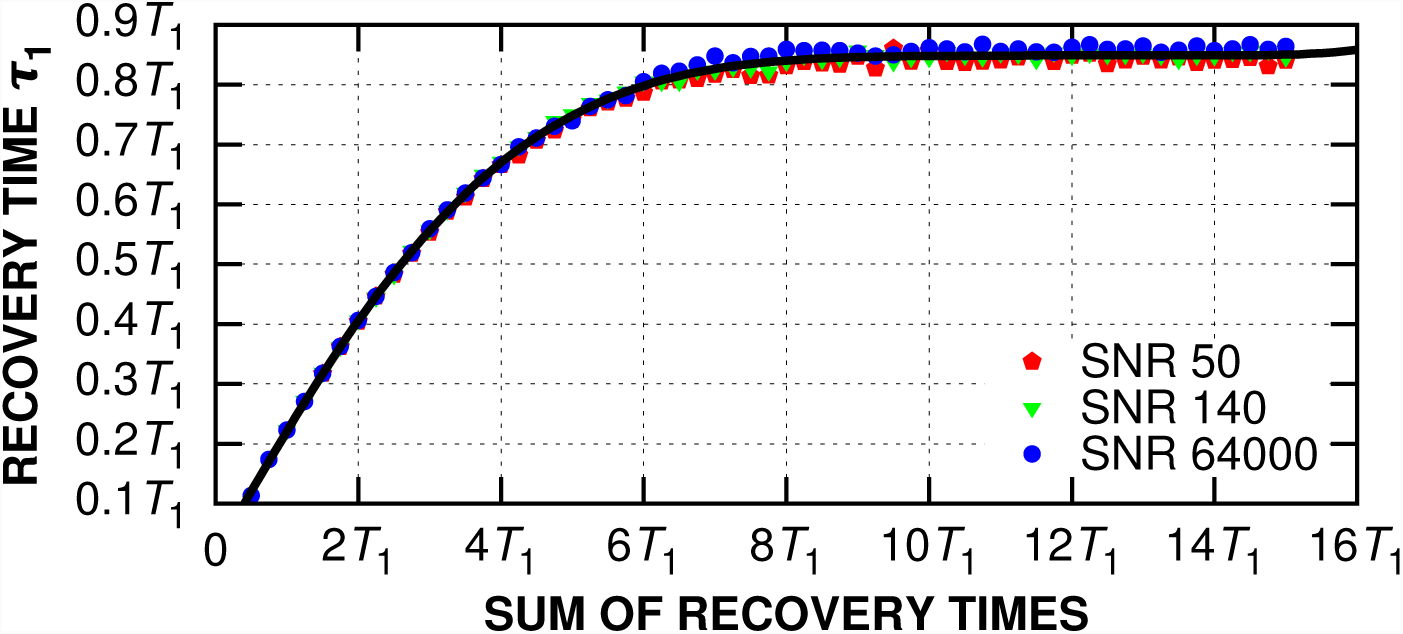
Optimal recovery time *τ*_1_ as a function of the sum of recovery times. The dots are the data points resulted from the computer simulation. The curve is described by Eq. (4) and is derived by least-squares fitting the data points.

where 0.05*T*_1_ ≤ *T*_Σ_ ≤ 15*T*_1_. This formula also independently reproduces the findings for the three cases considered in Ref. [8], in which the present Monte Carlo method was validated with the *R*_1_ measurement of brain CSF of human subjects. That is, the optimal *τ*_1_ for the SRT of 2*T*_1_, 3*T*_1_, and 6*T*_1_ was found to be 0.4*T*_1_, 0.55*T*_1_, and 0.8*T*_1_, respectively, whereas Eq. (4) produces 0.405*T*_1_, 0.556*T*_1_, and 0.799*T*_1_, respectively.

### Precision of the *R*_1_ Measurement

Figure 2 shows the standard deviation (SD) of the *R*_1_ distribution when the setting of the recovery times is optimal; that is, the figure shows the minimal SD that may be achieved for a given SRT *T*_Σ_ and SNR *ρ*, which can be described by

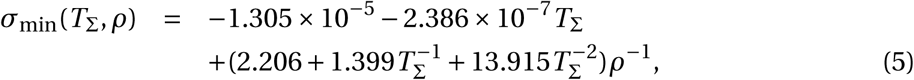

**Figure 2.**
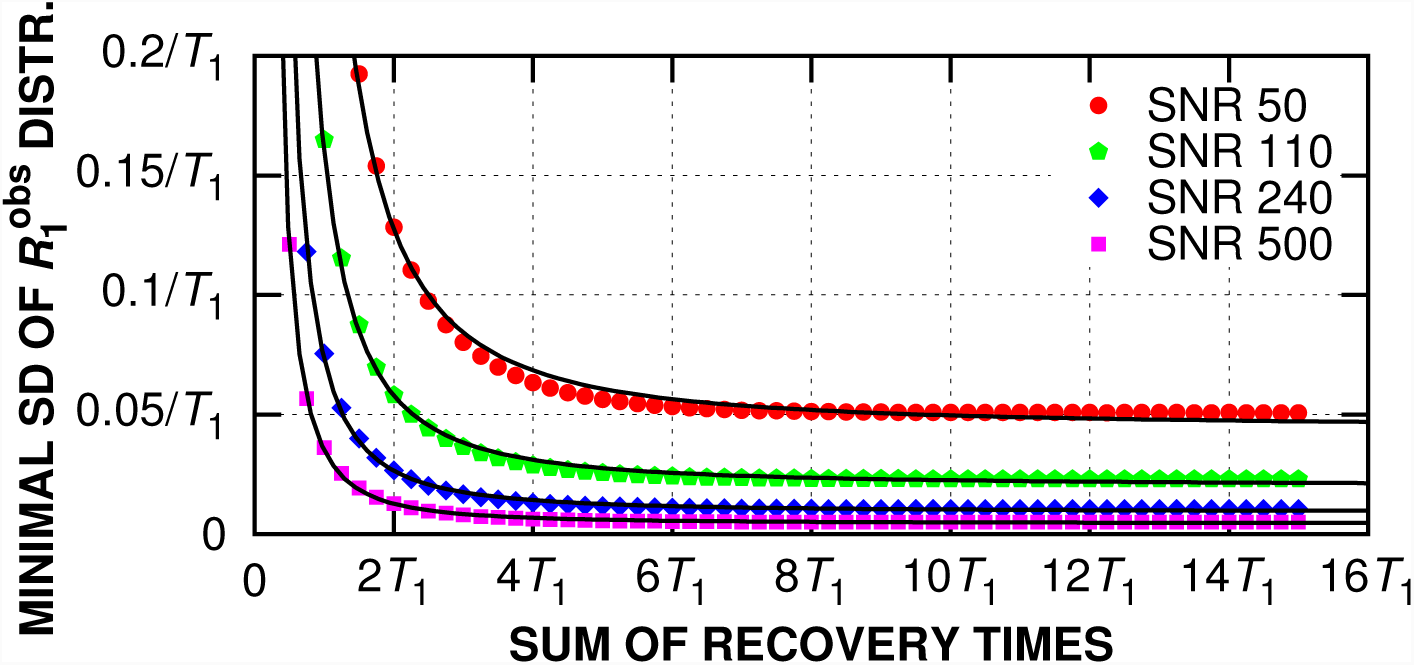
Minimal standard deviation (SD) of the computer-simulated 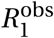 distribution as a function of the sum of the recovery times. The dots are the data points resulted from the computer simulation. The curves are described by Eq. (5) and are derived by least-squares fitting the data points.

for 0.05*T*_1_ ≤ *T*_Σ_ ≤ 15*T*_1_ and 50 ≤ *ρ* ≤ 64000. This formula provides a way to estimate the uncertainty of the *R*_1_ measurement. For example, because the standard error of the mean is related to the SD and the number of the measurements, with the SD calculated by Eq. (5) given the SRT and *ρ*, the number of measurements required to reach below the desired standard error can then be calculated. Similarly, for a given number of measurements and the SRT, the minimal SNR needed to achieve the desired standard error can be calculated by Eq. (5). An example is given in Ref. [8].

### Accuracy of the *R*_1_ Measurement

Figure 3 shows the mean of the computer-simulated 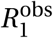 distribution that has the minimal SD. The effect of noise deviates the expected value of 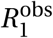 from the true *R*_1_ value towards a higher value, depending on the SNR and SRT. For the ranges of SNR and SRT studied in this work, the deviation is small. Compared with the magnitude of the SD (cf. Fig. 2), the deviation is negligible.

**Figure 3.**
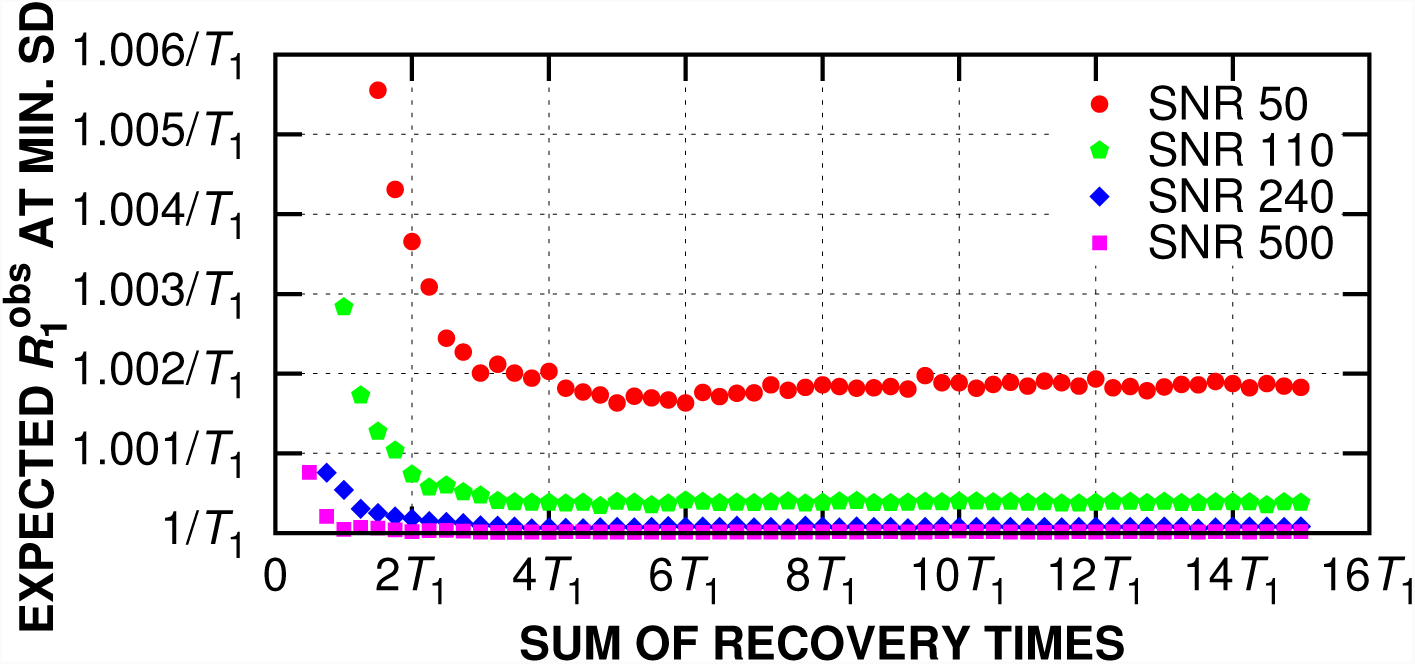
Mean of the computer-simulated 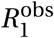 distribution when the selection of the recovery times produces the minimal standard deviation for the given sum of recovery times.

## DISCUSSION

As shown in Fig. 1, in the SNR range considered in this work, the minimal SD occurs at the same set of the recovery times despite the SNR. That is, the SNR is not a factor to consider in determining the recovery times, which is a property shared with the conditions in which the total scan time is not constrained and allows acquisition of more than two images [7, 9–11]. However, the SNR does affect the precision of the measurement, as shown in Fig. 2.

From Figs. 2 and 3, the precision and accuracy of the *R*_1_ measurement are rather sensitive to the SRT below ∼ 2*T*_1_, in which a small increase in SRT can produce a large improvement. However, increasing the SRT to improve the measurement is effective only up to a limit. For example, for SNR of 50, no more improvement can be gained when the scan time is extended beyond SRT of ∼ 6*T*_1_, and the strategy should be to increase the SNR.

In clinical settings, the determination of the SRT can depend on, for example, how long the scan subject can tolerate in the scanner and how many different types of scans (e.g., anatomical, diffusion, or functional) will be performed in a scan session. Because the SRT can influence the quality of the *R*_1_ measurement, a reference or a formula is needed to evaluate the precision and accuracy that can be achieved within the scan time allocated and to divide the scan time for the recovery times. Usually, the scan time can only allow acquisition of two images without signal averaging. In addition, given the same scan time, the improvement by using three points over two points is only marginal [8]. These are the conditions considered in the pulse sequence optimization in this work.

Optimization of the pulse sequence parameters has been of great interest in the development of quantitative NMR and MRI relaxation measurement (see, e.g., Kingsley [12] and references therein). The optimization typically seeks to find a configuration of the parameters such that the measurement outcome is least influenced by noise or random fluctuation. To include the scan time in the consideration, the precision per unit time (i.e., efficiency) is usually used as the target of optimization [13]; however, such optimization target implies the total scan time can be extended at will or the number of the points is variable [14], which could generate parameters that cannot be applied in the clinical settings described above. Therefore the constraint on the sum of recovery time is explicitly considered in this work, and the resultant Eq. (5) provides a way to evaluate the trade-offs between precision and the scan time.

Traditionally, equations are derived to describe the changes of the measurement outcome with respect to small changes of the parameters; then, the minima of the equations are located mathematically by, for example, differential calculus. In this work, the approach to optimization is different but shares a similar interpretation. The Monte Carlo method simulates the distribution of the measurement outcome under noise influence for a given configuration of the parameters and repeats the simulation for all reasonable configurations. Then the configuration that has the smallest standard deviation (SD) of the simulated distribution is considered optimal. As shown in Fig. 3, the optimal configurations also generate accurate means for the *R*_1_ measurement.

One limitation of the Monte Carlo method is that the simulation is performed for discrete samples of the parameters. To overcome this limitation, this work uses an interpolation function to explore the interval between samples as described in the Methods section. Another limitation has been the resolution power to distinguish small difference in the SD to locate the minimum, which is more important for the condition of a low SNR. For a low SNR, the SD is larger so the 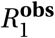 distribution is broader; thus a much larger number of trials would be needed to properly represent the distribution. If the number of trials is not sufficient, the simulated distribution is only a sample of the true distribution and the difference between the simulated and the true SD of the distribution can become appreciable and appear random, which can be seen in Fig. 1 in data of a larger SRT and in Fig. 3 in data of a lower SNR.

## CONCLUSION

Under noise influence, the accuracy and precision of the *R*_1_ measurement depend on the selection of the recovery times. In clinical application, the selection of the recovery times is subject to the constraint of the available scan time. This work, using an experimentally validated Monte Carlo computational method [8], provides general formulas describing the relationship between the total scan time, the optimal accuracy and precision, and the recovery-time selection for the two-point saturation–recovery method.

## References

[1] Weiss GH, Guptaj RK, Ferretti JA, Becker ED. The choice of optimal parameters for measurement of spin-lattice relaxation times. I. Mathematical formulation. J Magn Reson. 1980;37:369–379.

[2] Becker ED, Ferretti JA, Gupta RK, Weiss GH. The choice of optimal parameters for measurement of spin-lattice relaxation times. II. Comparison of saturation recovery, inversion recovery, and fast inversion recovery experiments. J Magn Reson. 1980;37:381–394.

[3] Hanssum H, Rüterjans H. Two-parameter least-squares analysis of inversion-recovery Fourier transform experiments. J Magn Reson. 1980;39:65–78.

[4] Kurland RJ. Strategies and tactics in NMR imaging relaxation time measurements. I. Minimizing relaxation time errors due to image noise–the ideal case. Magn Reson Med. 1985;2:136–158.

[5] Crawley AP, Henkelman RM. A comparison of one-shot and recovery methods in T1 imaging. Magn Reson Med. 1988;7:23–34.

[6] Doran SJ, Attard JJ, Roberts TPL, Adrian Carpenter T, Hall LD. Consideration of random errors in the quantitative imaging of NMR relaxation. J Magn Reson. 1992;100:101–122.

[7] Zhang Y, Yeung HN, O’Donnell M, Carson PL. Determination of sample time for T1 measurement. J Magn Reson Imaging. 1998;8:675–681.

[8] Hsu J-J, Glover GH, Zaharchuk G. Optimizing saturation–recovery measurements of the longitudinal relaxation rate under time constraints. Magn Reson Med. 2009;62:1202–1210.

[9] Jones JA, Hodgkinson P, Barker AL, Hore PJ. Optimal Sampling Strategies for the Measurement of Spin–Spin Relaxation Times. J Magn Reson B. 1996;113:25–34.

[10] Fleysher R, Fleysher L, Gonen O. The optimal MR acquisition strategy for exponential decay constants estimation. Magn Reson Imaging. 2008;26:433–435.

[11] Akçakaya M, Weingärtner S, Roujol S, Nezafat R. On the selection of sampling points for myocardial T_1_ mapping. Magn Reson Med. 2015;73:1741–1753.

[12] Kingsley PB. Methods of measuring spin-lattice (T1) relaxation times: An annotated bibliography. Concepts Magn Reson. 1999;11:243–276.

[13] Fleysher L, Fleysher R, Liu S, Zaaraoui W, Gonen O. Optimizing the precision-per-unit-time of quantitative MR metrics: Examples for T1, T2, and DTI. Magn Reson Med. 2007;57:380–387.

[14] Ogg RJ, Kingsley PB. Optimized precision of inversion-recovery T1 measurements for constrained scan time. Magn Reson Med. 2004;51:625–630.

